# The use of salivary α-amylase as an evolutionary solution to host selection in parasitoids

**DOI:** 10.1101/227173

**Authors:** Gladys Bichang’a, Jean-Luc Da Lage, Claire-Marie Mailhan, Frédéric Marion-Poll, Claire Capdevielle-Dulac, Michel Zivy, Thierry Balliau, Bruno Le Ru, Laure Kaiser-Arnauld, Gerald Juma, Esther Maina, Paul-andré Calatayud

## Abstract

Foraging parasitoids use chemical signals in host recognition and selection processes. Thereby, chemicals from the herbivore hosts play a crucial role. When different herbivores are present in the same plant or field, the perception of specific volatiles and contact compounds emitted from the host itself enable the parasitoids both to differentiate between hosts and non-hosts and to estimate the health status of its host. During the host feeding process, contact between the parasitoid and its host is very crucial, and oral secretions from the host play a key role during the first contact for such evaluation by the parasitoid. Using an integration of behavioral observations, biochemical and sensory physiological approaches we demonstrate that female parasitoids of *Cotesia flavipes* recognize their host and oviposit in reaction to an α-amylase, which is present in the oral secretions of the larvae of their host, *Chilo partellus*. This activity was also mediated by a purified α-amylase synthetized from *Drosophila melanogaster*. Using this synthetized enzyme, we further demonstrate that the conformation of the enzyme rather than its catalytic site is responsible for this activity. This enzyme is activating gustatory neurons of the terminal antennal sensilla chaetica of *C. flavipes* females. α-amylases are therefore good candidates for an evolutionary solution to host selection in parasitoids, thus opening new avenues for investigations in hosts-parasitoids interactions.

## INTRODUCTION

One of the strategies of biological control (BC) of pest insects is based on the use of natural enemies. Among natural enemies, insect parasitoids comprise the major biological control agents (Pimentel et al., 1992; Tilman et al., 2001; Lazarovitz et al., 2007; Godfray et al., 2010), able to control insect populations in the wild (Hawkins, 1994). Among insect parasitoids, *Cotesia* is one of the most diverse genera of the subfamily Microgastrinae (Hymenoptera, Braconidae), with almost 300 species already described (Yu et al., 2016) and probably over 1,000 species world-wide, e.g. Mason (1981). Many *Cotesia* species may appear generalists but careful ecological studies may reveal a hidden complexity with an assemblage of populations having a more restricted host ranges (Kaiser et al., 2017a).

In sub-Saharan Africa, lepidopteran stemborers of the Crambidae, Pyralidae and Noctuidae families are economically important pests of maize and sorghum (Harris, 1990; Polaszek, 1998; Kfir et al., 2002). Due to their widespread distribution and destructive nature, stemborers have been the subject of extensive research (Calatayud et al., 2006). The most cited species are the crambid *Chilo partellus* (Swinhoe), the noctuids *Busseola fusca* (Fuller) and *Sesamia calamistis* Hampson, and the pyralid *Eldana saccharina* (Walker)(Polaszek, 1998). With exception of *C. partellus*, which was accidentally introduced from Asia into Africa before the 1930s (Kfir, 1992), they are indigenous to Africa. During the early 1990s, the International Centre of Insect Physiology and Ecology (*icipe*) renewed emphasis on BC of *C. partellus* with the introduction of *Cotesia flavipes* Cameron (Hymenoptera: Braconidae) into Kenya from Asia. The parasitoid was first released in the coastal area in 1993 (Overholt et al., 1994), where it reduced *C. partellus* densities by over 50% (Zhou et al., 2001; Jiang et al., 2006). This was to complement the action of the closely related *Cotesia sesamiae* (Cameron) (Hymenoptera: Braconidae), which is the most abundant indigenous larval parasitoid of lepidopteran stemborers in ESA. However, parasitism by *C. sesamiae* is usually below 5% though in some localities it can attain 75% (Jiang et al., 2006; Kfir, 1995; Sallam et al., 1999; Songa et al., 2007).

The ability of parasitoids to successfully utilize cues in the two successive steps of habitat location, and discrimination between suitable and unsuitable hosts is crucial for the success of BC (Wajnberg et al., 2008; Wajnberg and Colazza, 2013). In the case of parasitoid targeting feeding host stage, the first step is often mediated by the volatile organic compounds (VOCs) resulting from the elicitation of plant defense metabolic pathways by salivary enzyme from the phytophagous host. When approaching the host, the parasitoids rely mostly on specific host-produces signals, and most of them are related to feeding activities, like fecal pellets and oral secretions (see Kaiser et al. [2017b] for a recent review).

Previous studies have shown that VOCs do not convey reliable information to *Cotesia flavipes* species complex, which includes *C. flavipes* and *C. sesamiae*, on the suitability of caterpillar species but they are mere indicators of the presence of herbivores (Ngi-Song and Overholt, 1997; Obonyo et al., 2008). It is only when approaching the host that reliable information on host’ identity is perceived for which tactile and contact-chemoreception stimuli from the hosts play a major role in host recognition and oviposition, and it is hypothesized that protein(s) present in the host’s oral secretions are involved (Obonyo et al., 2010a; 2010b; 2011).

In this study, an integration of behavioral observations, biochemical and sensory physiological approaches have been used to assess the nature of the active compound mediating host acceptance for oviposition, and to elucidate the mode of perception of this compound by the parasitoid, *C. flavipes*.

## MATERIALS AND METHODS

### Insects

*Cotesia flavipes* adults were obtained from laboratory-reared colonies established at the International Centre of Insect Physiology and Ecology (*icipe*), Nairobi, Kenya. The colony originated from individuals collected in the field in the coastal region of Kenya in 1998. Field collected *C. flavipes* were added twice a year to regenerate the colony. The parasitoid was reared on *C. partellus* larvae according to the method described by Overholt et al. (1994). Parasitoid cocoons were kept in a Perspex cage (30 cm x 30 cm x 30 cm) until emergence. Adult parasitoids were fed on a 20% honey/water solution presented. They were then put under artificial light and left for 24 h to mate. In all the behavioral bioassays, only 1-day-old naïve, mated females were used. Experimental conditions were maintained at 25 ± 2 °C, 50–80% relative humidity (RH), and a 12:12 h (L:D) photoperiod (Overholt et al., 1994).

The host *C. partellus* originated from maize grown in the coastal region of Kenya. The larvae were reared on the artificial diets described by Ochieng et al. (1985). Thrice a year feral stemborer larvae from the coastal region were added to rejuvenate the colonies.

### Collection of oral secretions from *Chilo partellus* larvae

It is known that acceptance of *C. partellus* host larvae for oviposition by *C. flavipes* is enhanced when the host larvae were fed on maize stems for 24h prior exposure to parasitism (Mohyuddin *et al*. 1981; Inayatullah, 1983; Van Leerdam *et al*., 1985; Potting *et al*., 1993; Overholt *et al*. 1994). Therefore, to isolate the semiochemicals of the oral secretions of *C. partellus* that can be involved in host acceptance of *C. flavipes*, we used larvae previously fed for 24h on their original host plant (maize stems) and also, for comparisons, on stems of an alternative host, *Penisetum purpureum* Schumach. (Poaceae), surrounding frequently maize farms in Kenya. We compared also the behaviour of *C. flavipes* towards these two types of oral secretions with oral secretions of larvae fed on artificial diet of Ochieng *et al*. (1985). In addition, to verify if these semiochemicals are synthesized when the host are feeding. We compared also the oral secretions from starved larvae for 48h. For each type of oral secretions collection, a single larva held by a soft forceps was squeezed behind the head and capillary tube was used to collect oral secretions and placed directly on ice. The process was repeated for several larvae. The volume of oral secretion was estimated by weighting. All samples were preserved at -80°C before use. As evoked at the introduction, in a previous study it was hypothesized that the semiochemicals from oral secretions involved in host recognition by *C. flavipes* might include enzymes or thermo-labile proteins (Obonyo *et al*. 2010b). Therefore, we compared also oral secretions from larvae fed on maize stems but previously treated by proteinase K (Sigma product P6556) in order to destroy the proteins present in the oral secretions. In summary, the following types of oral secretions were compared:

- from starved larvae;
- from larvae fed on maize stems;
- from larvae fed on *P. purpureum* stems;
- from larvae fed on artificial diet;
- from larvae fed on maize stems followed by proteinase K digestion.

### Behavioral bioassays

In previous studies, we demonstrate that the parasitic wasps exhibit antennation (=use of antennae to prospect by drumming the body of the host) followed by at least one stinging attempt (one tentative of insertion of their ovipositors in the host) to accept a caterpillar as a host for oviposition (Obonyo et al., 2010a; 2010b). Therefore, in this study we used these behavioral steps to evidence host acceptance by *C. flavipes*. To test the behavioural activities triggered by different extracts (i.e. type of oral secretions, electrophoretic bands and known proteins, see previous and next sections), they were placed on small cotton wool presented to female wasps. A small piece of cotton wool was rolled into spherical shape (around 2 mm in diameter) and placed at the centre of a Petri dish of 8 cm diameter without the Petri dish cover. About 0.5 to 1 μl of the extract to be tested was deposited on the cotton wool ball while ensuring that the cotton wool was kept moist but not wet. A single female wasp was introduced near the cotton wool and both were covered with a transparent circular Perpex lid (3 cm diameter, 1 cm height) to prevent the parasitoid from flying off and to allow the observations.

The behaviour of the wasp in the Petri dish was then monitored for a maximum of 120 s. For each wasp, both the antennation and stinging attempt were recorded. The percentage of positive response (i.e. antennation + stinging) was calculated from 10, 20 or 30 wasps tested per electrophoretic bands, per type of oral secretions or per identified proteins (see previous and next sections), respectively. The wasp, the cotton wool ball with tested extract and the arena were replaced each time between each observation.

All behavioural experiments were carried out in a room with temperature of 26 ± 1 °C between 10h00 to 14h00 with a constant source of light to maintain an optimal temperature for the behavioural activities of the female wasps.

### Electrophoresis and isolation of proteins from polyacrylamide gel

The oral secretions from *C. partellus* were first centrifuged at a maximum speed of 14,000 ×g for 5 minutes in order to remove the undetected debris (frass and undigested food materials). This was followed by desalting and concentrating the samples using Amicon^®^ Ultra-0.5 centrifugal filter devices (Merck Millipore). The samples were quantified before electrophoresis using the Pierce BCA protein assay Kit (Thermo Scientific No. 23227) based on bicinchoninic acid (Smith et al., 1985). All the quantification measurements were carried out using Eppendorf-Biospectrometer fluorescence machine (SN 667).

Electrophoresis was conducted under non-denaturing conditions (native PAGE electrophoresis, Ornstein-Davis discontinuous buffer system) according to the method described by Chrambach and Jovin (1983) and Niepmann and Zheng (2006). The gels were cast in two sections using the Bio Rad Mini-PROTEAN^®^ Electrophoresis System and Hoefer™ Mini Vertical Electrophoresis Systems (Fisher Sci.com). A stacking gel (4%T, 2.7%C, 0.125M Tris-Cl pH 6.8) was cast on top of a resolving gel of (7.5%, T4.4%C, 0.125M Tris-Cl pH 6.8). Electrophoresis was conducted (running buffer: 0.025M Tris, 0.192M glycine pH 8.3) immediately after loading the samples at a constant voltage of 150V and current of 25mA for 1-2hr in a cold room. At the end of the run, gels were immediately removed and stained for 30 min in a staining solution consisting of 0.2 % Coomassie Brilliant Blue R250. The gels were then destained with a solution of methanol, glacial acetic acid and water at the ratio of 4:1:5. The stained proteins were compared with a molecular mass standard (Sigma Aldrich) containing albumin from bovine serum (Sigma A8654, 132 kDa), urease from jack bean (Sigma U7752, 272 and 545 kDa), α lactalbumin from bovine milk (Sigma L4385, 14.2 kDa) and albumin from chicken egg white (Sigma A8529, 45 kDa).

For the isolation of electrophoretic bands, the protein bands were manually excised from the gel before staining process following the method of Kurien and Scofield (2012) with some modifications. The excised gel fragments containing the protein of interest were frozen overnight at -80°C. Each frozen gel fragment was ground using a mortar into fine powder under liquid nitrogen and the resulting gel powder transferred to the upper chamber of the Costar^®^ column (centrifuge tube filters, Costar lot No. 22304012 Corning incorporated, NY 14831-USA). The protein trapped in the gel powder was eluted using native elution buffer 0.25M Tris HCl buffer pH 6.8, or normal saline depending on the subsequent application. After 10 min of centrifugation at 13000 ×g, 300 to 350 μl of the filtrate was recovered and stored for further concentration and desalting. A second elution was performed with fresh elution buffer and a filtrate of approximately 250-300 μl was collected and combined with the previous one. Each protein eluted was concentrated 25-30 × folds using a Amicon centrifugal device equipped with 30K MWCO omega membrane. The concentrated protein eluents were assayed for protein content with the aforementioned Pierce BCA protein assay Kit. For each protein eluent, the purity and elution efficiency were checked by native PAGE electrophoresis. Proteins in the gel were Coomassie-stained as above. All the 7 major bands revealed in the oral secretion of *C. partellus* fed on maize (see Fig. 1) were separated and purified as described above for use in behavioural assays (see previous section).

**Fig. 1.**
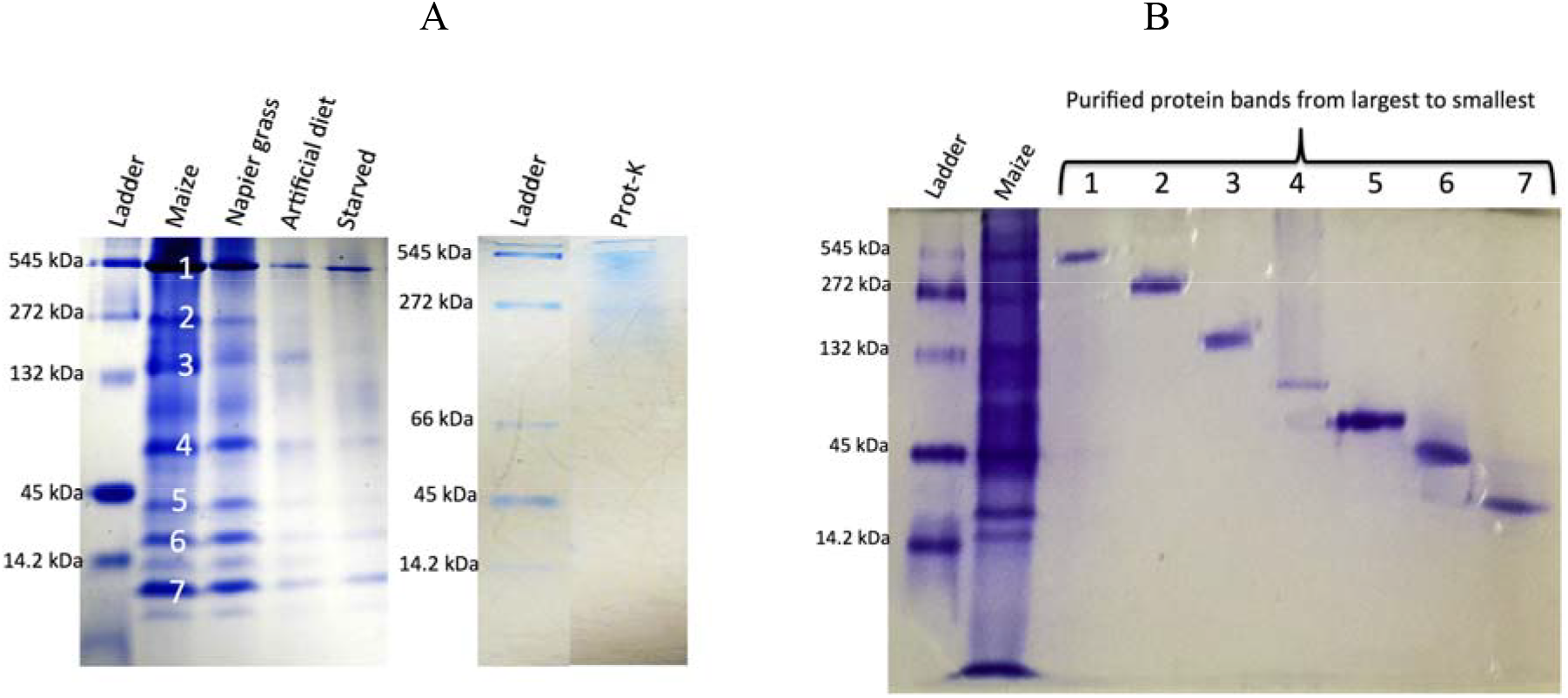
Analysis of oral extracts in a native gel system. Protein samples were separated by 1D gel, 7% native Onstein-Davis discontinuous (Tris-glycine) PAGE before Coomassie staining. A) Comparison of *Chilo partellus* oral extract fed on different diet. Ladder: Sigma molecular weight markers; lane 1: oral secretion from *Chilo partellus* larvae fed on maize stems (Maize)(each main electrophoretic band [noted 1 to 7 on the gel] were individually extracted from the gel (see Fig. 1B) under non-denaturing conditions and tested towards *Cotesia flavipes* (see Table 2); lane 2: oral secretion from *Chilo partellus* larvae fed on *Pennisetum purpureum* stems (Napier grass); lane 3: oral secretion from *Chilo partellus* larvae fed on artificial diet (Artificial diet); lane 4: oral secretion from starved larvae of *Chilo partellus* (Starved). For each lane, 15μl of the oral secretion was loaded after concentrating and before quantification of the samples (Bio Rad Mini-PROTEAN^®^ Electrophoresis System). After proteinase K treatment no band was obtained (Prot-K). B) Individual protein band purified from the gel of regurgitant of *Chilo partellus* fed on maize. Lanes: 1 molecular weight marker (sigma Aldrich), 2 regurgitants from *Chilo partellus* fed on maize (Maize); lanes 1-7 bands purified and tested for activity against *Cotesia flavipes* (Hoefer^™^ Mini Vertical Electrophoresis Systems (Fisher Sci.com) (see Table 2).

### Protein identification

The gel purified protein eluent inducing parasitoids’ host recognition and oviposition were identified using LC-MS/MS. The protein eluent were first denatured in Laemmli buffer and then concentrated using a short electrophoretic migration, which also allowed removing any contaminants that could interfere with the trypsic digestion. Electrophoretic bands were excised and the gel pieces were washed in successive baths consisting of 50mM ammonium bicarbonate and acetonitrile. Proteins were then reduced by 10 mM of 1.4 dithiothreitol (DDT) and alkylated with 55mM of iodoacetamide to block the sulfide bonds of cysteines. After rinsing to remove residues of DTT and iodoacetamide, proteins were hydrolyzed by the addition of 0.125μg trypsin for 7 hours. After hydrolysis, the resulting peptides were extracted from the gel pieces with 50% acetonitrile acidified with 0.5% of trifluoroacetic acid (TFA). After complete speed vac drying, peptides were resuspended in a solution of 2% acetonitrile, 0.05% formic acid and 0.05% trifluoroacetic acid. Peptide mixes were then analyzed by LC-MS/MS using a nanoRSCL (thermoFinnigan) coupled with a LTQ Orbitrap Discovery (Thermo). The samples were loaded on a PepMap100C18 trap column for 5min with 2% acetonitrile (ACN), 0.08% TFA qsp H2O. Two buffers systems were used to elute the peptides: 2%ACN and 0.1% formic acid in water (buffer A); 98% ACN and 0.1%formic acid in water (buffer B). Peptide separation was performed using a linear gradient from 4% to 38 % of buffer B in 15min. The nanoHPLC was connected to the mass spectrometer using a nano electrospray interface (non-coated capillary probe 10μ I.d. New objective). Peptides ions were analyzed using Thermo Xcalibur (version 2.0.7) using the following data dependant steps: (1) full MS scan with a 300 to 1400 m/z range in the Orbitrap with a resolution of 15,000; (2) fragmentation by CID in the linear trap with a normalized energy at 35%. Step 2 was repeated for the three most intense ions with a minimum intensity of 500. Dynamic exclusion was set to 30 seconds.

Raw files were converted to the mzxml format using msconvert (3.0.9576 <u>http://proteowizard.sourceforge.net/tools.shtml</u>). Database search was performed using X!tandem JACKHAMMER (Craig and Beavis, 2004). Tolerance was set to 10 ppm for precursor ions and 0.5 Th for fragment ions. Cys-carboxyamidomethylation was set to static modification. Methionine oxydation, Nter acetylation of proteins, glutamine Nter deamidation and glutamic acid Nter water loss were set to variable modifications. Three databases were used: the *Spodoptera frugiperda* (Smith) EST database (http://www.ncbi.nlm.nih.gov/nucest version 2015, translated in the six reading frames and filtered to a minimum of 80 amino acids; 392,538 entries); the *Zea mays* database (from maizegdb, version v5a; 136,770 entries) and a standard contaminant database (55 entries). Identified peptides were filtered using X!tandemPipeline v3.3.4 (Langella et al., 2016) with the following criteria: peptide E-value less than 0.03, minimum 2 peptides per protein, protein E-value less than 10^−4^. Unassigned spectra were subjected to *de novo* identification using denovopipeline v1.5.1 (<u>http://pappso.inra.fr/bioinfo/denovopipeline/</u>), that allows the selection of unassigned spectra of good quality and their submission to pepnovo (v2010117, Frank 2005). Spectrum quality score was set to 0.2 and pepnovo score to 70. *De novo* sequences were then aligned to the same databases as for X!Tandem search using Fasts.v36.06 (Mackey et al., 2002). Proteins with a homology score less than 10^−4^ were validated. The biological and analytical reproducibility were addressed by a quantitative western blot (see next section).

Identified EST sequences obtained from digested peptides were submitted to a BLAST procedure (BLASTX, NCBI). The resulting protein was characterized by the name, the source and the molecular weight and a E-value/log E-value coverage. In order to calculate the coverage per cent of a peptide, the EST sequence was translated into a protein sequence using the Expasy Translate tool (http://www.expasy.org/tools/dna.html).

### Western blot analysis of the protein eluent inducing parasitoid oviposition

In order to confirm that the proteins purified and identified were indeed α-amylases, we performed a western blot using an antibody specific to *Drosophila melanogaster* Meigen α-amylase. Ten microliters of each heat denatured protein sample (of about 500 ng/μl) were loaded on a NuPAGE 4-12% Bis-Tris Gel (Invitrogen) and electrophoresis conducted for one hour at 200 volt in MOPS buffer. The proteins were then transferred to an iBlot Gel Transfer Nitrocellulose membrane (Invitrogen) using the iBlot Gel Transfer Device (Invitrogen). The membrane was washed in 1X PBS for 20 minutes, after which it was incubated for 90 minutes in a milk solution (1X PBS, 0.1% Tween, 5% milk) in order to saturate the membrane with proteins. The membrane was then incubated with the primary anti *Drosophila melanogaster* α-amylase antibody, kind gift from Dr B. Lemaitre (Chng et al., 2014), 1000-fold diluted in a solution of 1X PBS, 0.1% Tween, 1% milk) for several hours. After this step, the membrane was washed six times in 1X PBS, 0.1% Tween before incubating with the secondary antibody (Anti guinea pig IgG Peroxidase, Sigma A7289), 1000-fold diluted in a solution of 1X PBS, 0.1% Tween, 1% milk, for one hour The membrane was then washed 3 times in 1X PBS, 0.1% Tween. The peroxidase activity was detected with Amersham ECL Prime Western Blotting Detection Reagent (GE Healthcare) and recorded on an Odyssey FC imager.

### Sources of different α-amylases assayed

To confirm the involvement of α-amylases in host acceptance and oviposition by *C. flavipes*, we used well-purified and well-identified α-amylases from different organisms available in the commerce or in our lab. at Gif-sur-Yvette: the micro-organism, *Aspergillus oryzae* (Ahlburg) E. Cohn, the insects, *Drosophila melanogaster* and *Chilo suppressalis* (Walker); and the pig as a mammal (porcine pancreas). A-amylases from *A. oryzae* and porcine pancreas were obtained from Sigma No A9857 and A3176, respectively. The α-amylase from *Drosophila melanogaster* was produced in the yeast *Pichia pastoris* (Guillierm) Phaff, as described in Commin et al. (2013). The α-amylase of *C. suppressalis* was also produced in *P. pastoris*: the coding sequence of the *C. suppressalis* amylase gene 108827 was synthetized (Eurofins MWG), with replacement of the signal peptide by the one of *D. melanogaster* amylase (suppl. Fig. S1). We assayed an amylase from *C. suppressalis*, because its genome is available, contrary to *C. partellus*. In addition, to check if the behavioural activities of *C. flavipes* triggered by α-amylase (see results) was due to the structural conformation and/or the catalytic activity. We synthesized an inactive α-amylase with no change in its structural conformation. An inactivated α-amylase of *D. melanogaster* was obtained by a single replacement of the crucial catalytic residue Asp186 by an asparagine, which does not change the structural conformation (Aghajari et al., 2002). A colorimetric activity test (Infinity Amylase Reagent, Thermo Fisher) was used to confirm that this α-amylase of *D. melanogaster* had no catalytic activity.

### Electrophysiological responses from wasp antennal sensilla towards α-amylases

Similarly to Iacovone et al. (2016), electrical activity was recorded from antennal sensilla chaetica of the female wasp in response to the protein extract and reference compounds using the tip-recording technique (Hodgson et al., 1955). Female wasps (1–3 days old) were secured to a platform using thin strips of adhesive tape. The insect was grounded via a silver wire, bridged to the insect body by a drop of electrolyte gel (Redux^®^ Gel, Parker laboratories, Inc. Fairfield, NJ). Individual sensilla chaetica were contacted at the tip with a glass electrode containing the taste solution and an electrolyte (tricholine citrate 30 mM) which ensures a good electrical contact as well as inhibits the gustatory neuron to water and elicits not more than 8 spikes/s (Fig. 5). In Drosophila, tricholine citrate ensures a good electrical contact and inhibits the water cell (Wieczorek and Wolff, 1989). Taste responses were recorded for 2 s and were performed under a microscope (Z16 Apo, Leica France). Electrodes (borosilicate glass capillaries, 1.0 mm O. D. x 0.78 mm I. D., Harvard Apparatus) with a tip diameter of approximately 10 μm were pulled using a laser electrode puller (Model P-2000, Sutter Instrument Co, USA).

The recording electrode was connected to a preamplifier (gain = x10; TastePROBE DTP-02, Syntech, Hilversum, The Netherlands) (Marion-Poll and van der Pers, 1996), and the electric signals were further amplified and filtered by a second amplifier (Cyber-Amp 320, Axon Instrument, Inc., gain = x100, eight-order Bessel pass-band filter = 10–2800 Hz). These signals were digitized (DT9818, Data Translation; sampling rate = 10 kHz, 16 bits), stored on computer, and analysed using dbWave (Marion-Poll, 1996). Spikes were detected and analysed using software interactive procedures of dbWave. We evaluated the action potential frequency by counting all spikes occurring during the first second of recording.

The responses to the following stimulants were recorded extracellularly and compared using 30 mM tricholine citrate (all compound tested were suspended into this solution to inhibit the gustatory neuron to water) as control:

- oral secretion of *C. partellus* as a control;
- purified α-amylase from *D. melanogaster* and *C. suppressalis* at 300 ng/μl (the concentration of the band n°4 that was inducing oviposition in *C. flavipes*, see results);
- BSA at 300 ng/μl (as a standard protein of 55-60 kD, molecular weight close to α-amylase).

### Statistical analysis

The Marascuilo’s procedure was used to separate the percentages of wasps that exhibited positive responses (i.e. antennation + stinging attempts) (Marascuilo, 1966). For bioassays with known proteins, the percentage of positive response was calculated from a group of 5 wasps replicated 6 times (i.e. n=6). A non-parametric Kruskal-Wallis test was applied with type of proteins as factor. ANOVA was not used because none of the data were normally distributed and had homoscedastic variance. Following Kruskal-Wallis test, a pairwise Wilcoxon’s rank sum test was conducted with false discovery rate (FDR) correction for multiple testing. Comparisons among sensilla chaetica responses towards oral secretions of *Chilo partellus* and different proteins were conducted using one-way analysis of variance (ANOVA) with the Tukey’s contrast test for multiple comparisons between means. Before running this ANOVA, the homogeneity of variance and data normality were examined by F - test and Kolmogorov–Smirnov methods. These statistical analyses were done in R version 3.3.1 (2016).

## RESULTS

The oral secretions of *C. partellus* previously fed on maize stems induced significant antennation and stinging attempt (Table 1). The *C. partellus* oral secretions from larvae previously fed on *P. purpureum* triggered as many responses as the one from maize-fed host larvae. Comparatively, oral secretions of larvae fed on artificial diet did not elicit any behavioral activity. In addition, the oral secretions from larvae starved for 48h did not elicit any behavioral response as well as when the oral secretions from larvae fed on maize stems were treated with proteinase K.

**Table 1.**
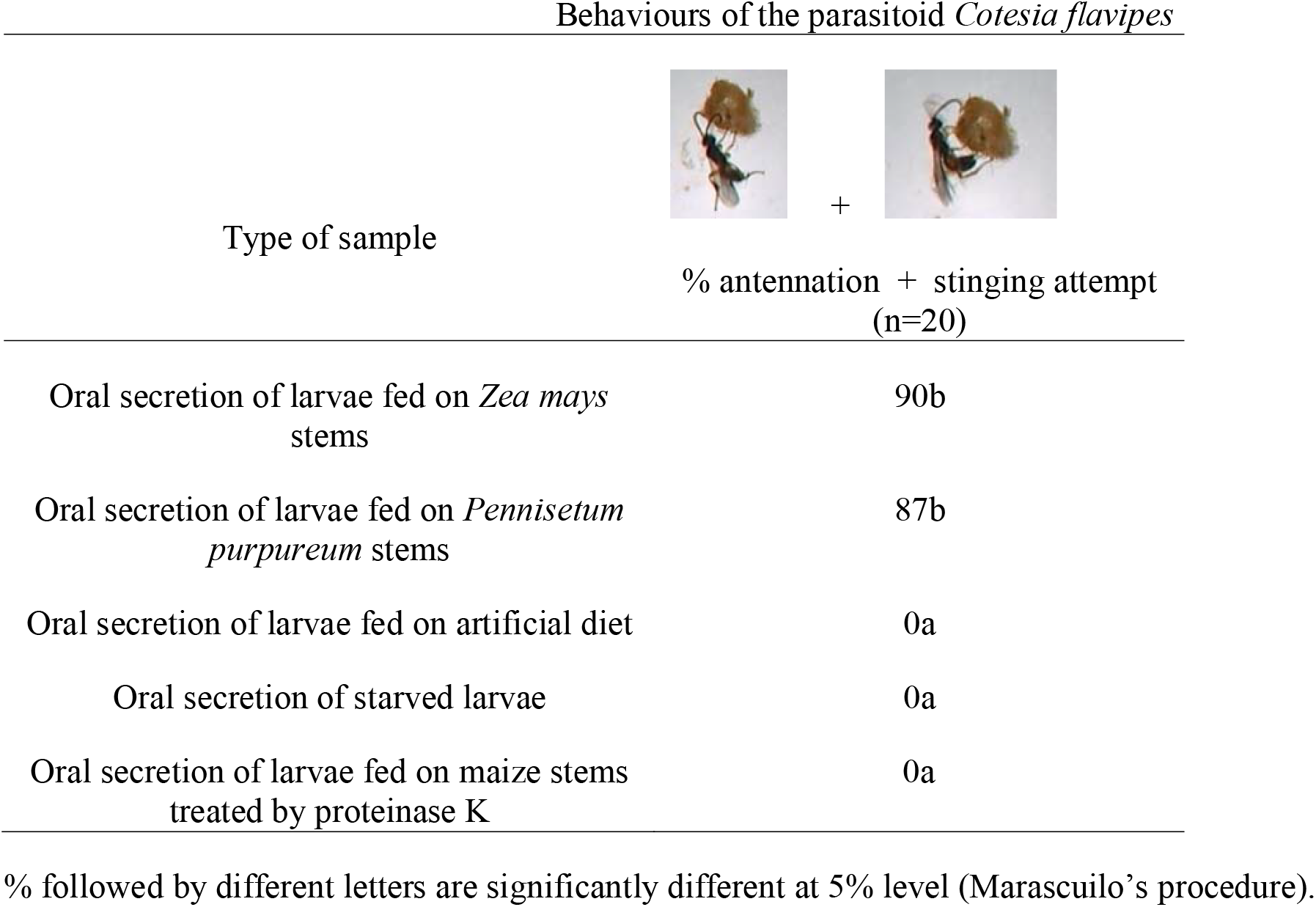
Response of *Cotesia flavipes* parasitic wasps to oral secretions of its host, larva of *Chilo partellus*.

Behaviours of the parasitoid *Cotesia flavipes*
| Type of sample |  |
| --- | --- |
|  | % antennation + stinging attempt<br>(n=20) |
| Oral secretion of larvae fed on *Zea mays* stems | 90b |
| Oral secretion of larvae fed on *Pennisetum purpureum* stems | 87b |
| Oral secretion of larvae fed on artificial diet | 0a |
| Oral secretion of starved larvae | 0a |
| Oral secretion of larvae fed on maize stems treated by proteinase K | 0a |

% followed by different letters are significantly different at 5% level (Marascuilo's procedure).

The electrophoretic analyses of the active oral secretions revealed the presence of more intense electrophoretic bands (i.e. higher quantities of proteins) than of the inactive oral secretions, confirming the involvement of protein(s) in triggering antennation and stinging attempt (Fig. 1A).

The oral secretion of larvae fed on maize stems showed seven major electrophoretic bands in a one-dimension gel electrophoresis under non-denaturing conditions (Fig. 1A). Each major band was manually excised from the gel, extracted (Fig. 1B) and tested for further behavioral responses as shown in Table 2. Out of these seven protein bands, only two bands elicited activity, particularly band no 4 (≈ 50 kDa) which triggered the highest response, i.e 90% of *C. flavipes* exhibited antennation and stinging attempt (Table 2). It was thus subjected to further analysis and identification.

**Table 2.**
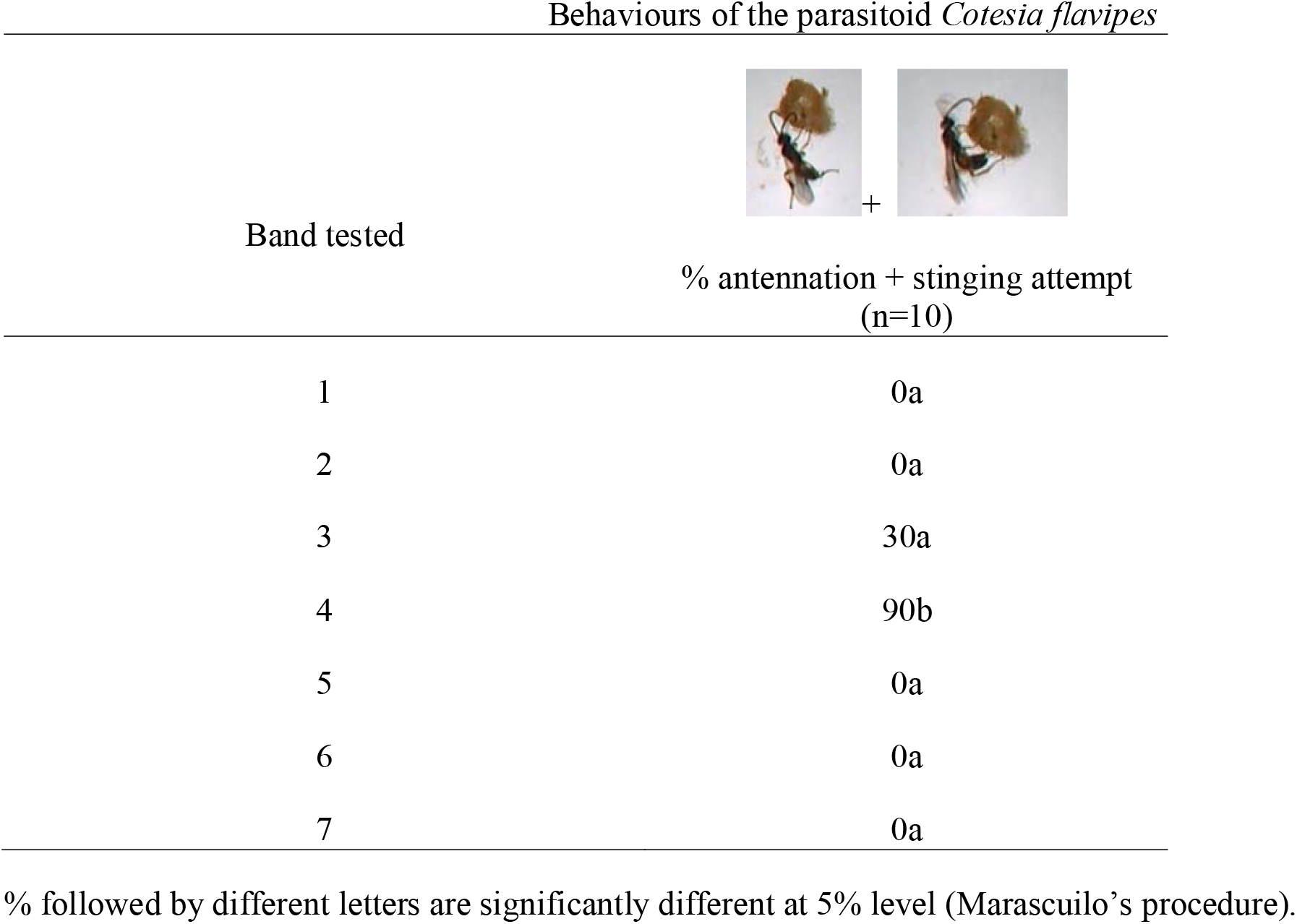
Response of *Cotesia flavipes* parasitic wasps to the seven main electrophoretic bands (see Figure 1) obtained from the oral secretions of its host, larva of *Chilo partellus*.

| Behaviours of the parasitoid *Cotesia flavipes* |  |
| --- | --- |
| Band tested | % antennation + stinging attempt<br>(n=10) |
| 1 | 0a |
| 2 | 0a |
| 3 | 30a |
| 4 | 90b |
| 5 | 0a |
| 6 | 0a |
| 7 | 0a |

% followed by different letters are significantly different at 5% level (Marascuilo's procedure).

In order to identify the active protein band that induced the highest behavioral response, proteins from band No 4 were digested and the resulting peptide mixture was analyzed by liquid chromatography-mass spectrometry. Database search allowed the identification of two distinct maize proteins with 5 and 2 peptide sequences, respectively, while *de novo* sequencing allowed the identification of 22 peptides that matched to accession gi|295290041|gb|FF379314.1|FP379314| of the *S. frugiperda* database of mid gut cDNA sequences (Supplementary Table 1). The protein sequence blasted significantly with α-amylase superfamilies (Fig. 2). The confirmation of α-amylase assignation of the electrophoretic band no 4 was done by western blot analysis (Fig. 3). The anti-α-amylase of *D. melanogaster* linked mostly with the band no 4 (≈ 50 kDa) of the oral secretion of *C. partellus* and with that extracted from the gel.

**Fig. 2.**
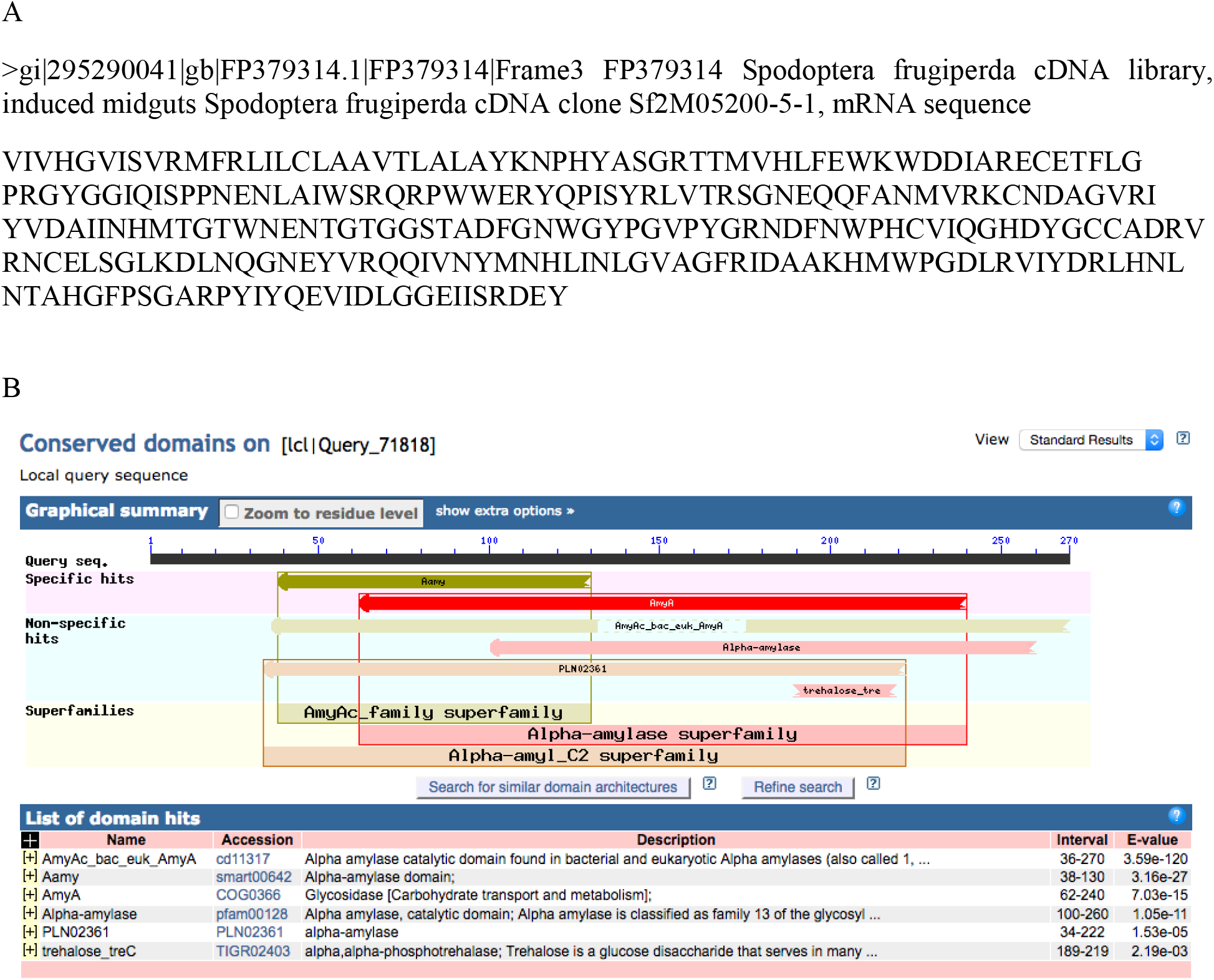
Protein-protein BLAST result of the *de novo* protein sequence. A) The best *de novo* protein sequence associated with EST specific to *Spodoptera frugiperda* database (see Table S1). B) The precomputed domain annotation for the best *de novo* protein sequence of A) using the protein data bank of the BLAST ® online software (<u>https://blast.ncbi.nlm.nih.gov/Blast.cgi</u>). The section circled in red provides the functional label that has been assigned to the subfamily domain.

**Fig. 3.**
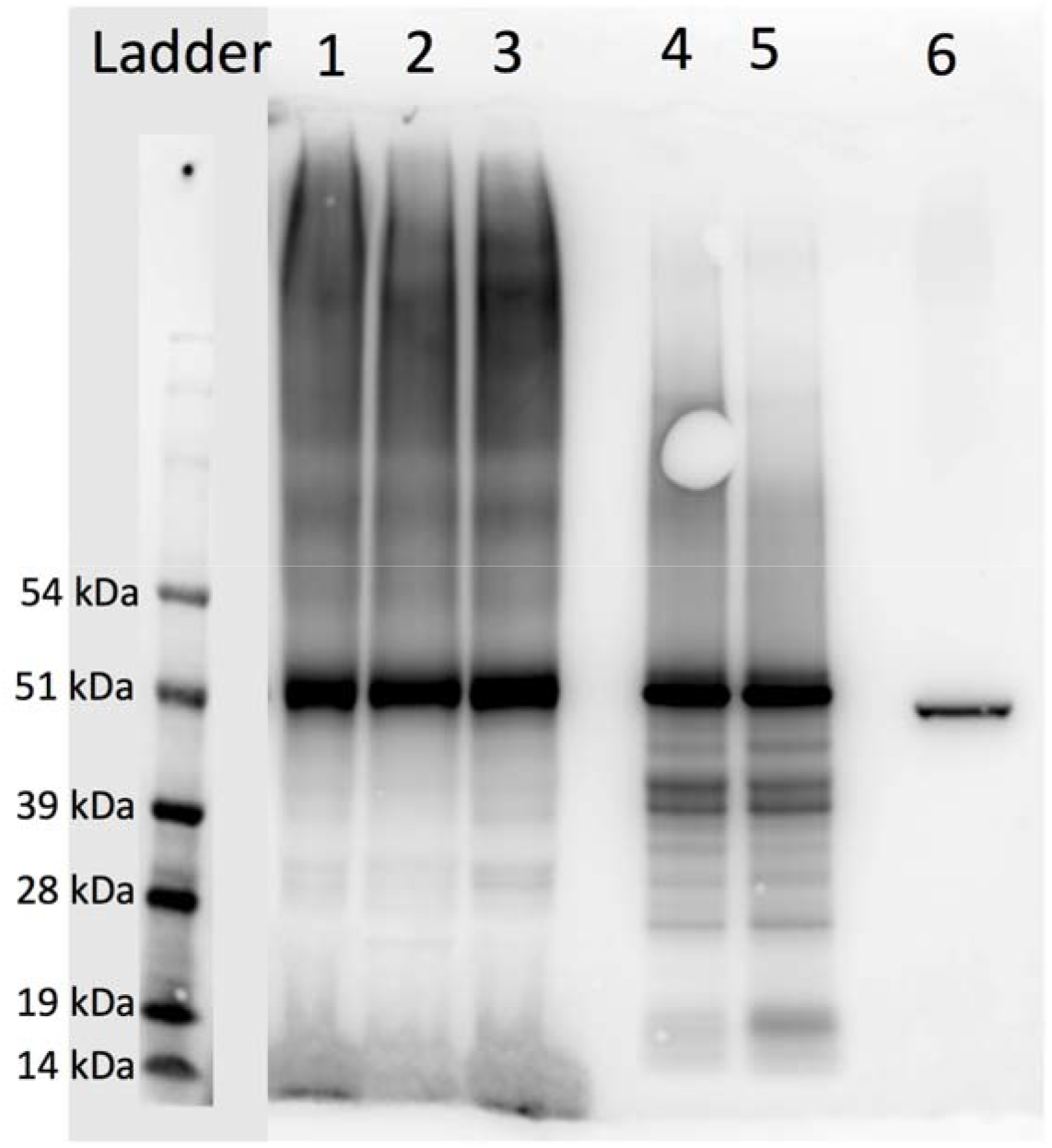
Western blot performed with an antibody specific to *Drosophila melanogaster* α-amylase. Ladder: molecular weight markers (pre-stained SeeBlue Plus2, Thermo Fischer); 1, 2 and 3: oral secretions from *Chilo partellus* larvae fed on maize stems; 4 and 5: band n°4 of Fig. 1 which has been extracted from the gel and used for Western Blot analysis; 6: α-amylase from *Drosophila melanogaster*.

The activity elicited by different α-amylases from different origin, including α-amylase of *D. melanogaster*, confirmed the involvement of this enzyme in *C. flavipes* antennation and stinging attempt (Table 3). In contrast, the use of a different protein such as BSA did not induce any behavioral response in the wasp. The α-amylases from insects, i.e. *D. melanogaster* and *C. suppressalis*, induced the highest behavioral responses in *C. flavipes* antennation and stinging attempt although not significantly different to the responses induced by *A. oryzae* α-amylases (Table 3). To check if the behavioral activity of *C. flavipes* triggered by α-amylase was due to the structural conformation and/or the catalytic activity, we used an inactivated α-amylase from *D. melanogaster* with no change in its structural conformation. Interestingly this inactivated α-amylase still induced behavioral activities of *C. flavipes* indicating that the conformation rather than the catalytic activity of α-amylase is crucial in the host acceptance process by *C. flavipes*.

**Table 3.**
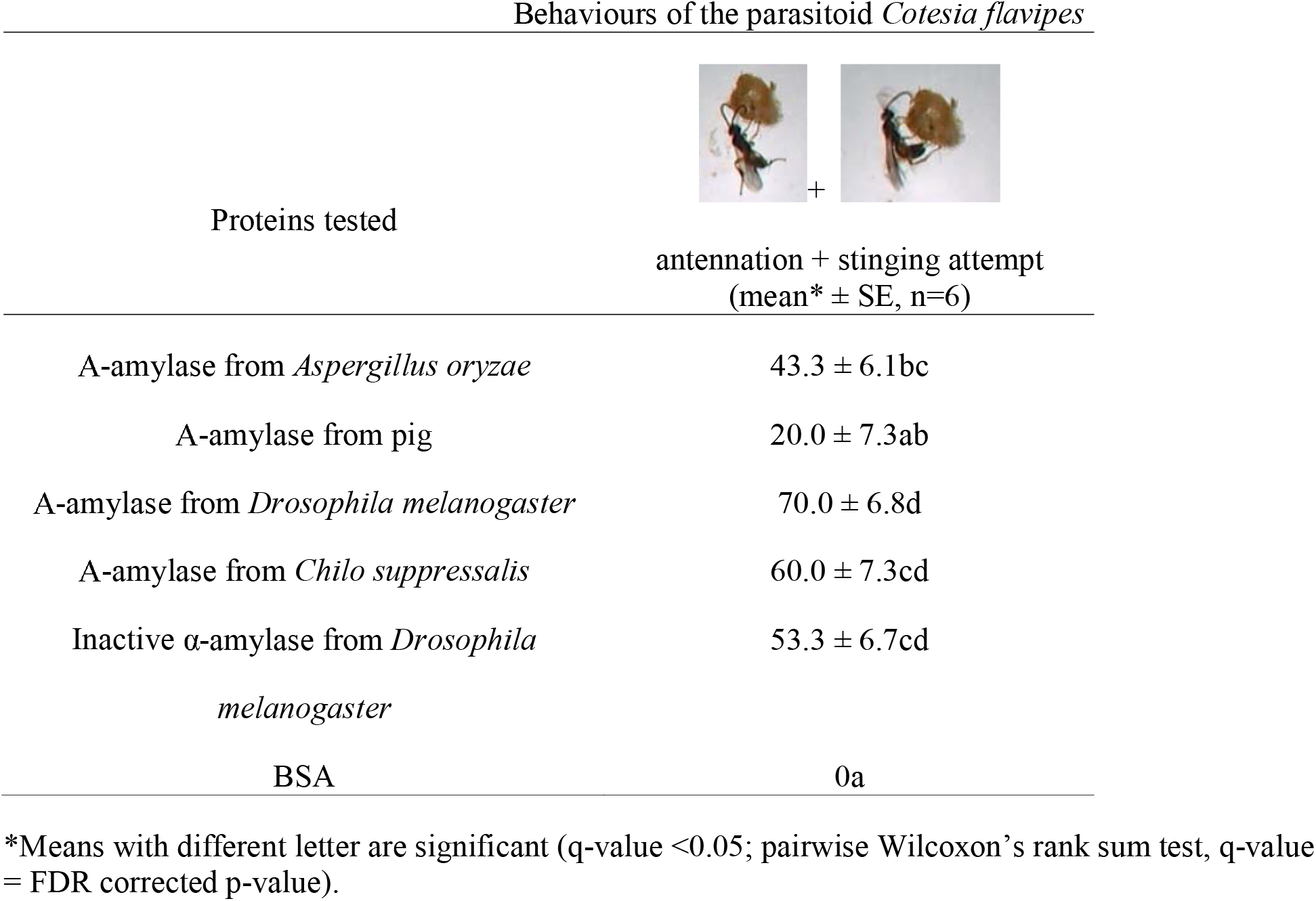
Response of *Cotesia flavipes* parasitic wasps to purified proteins (at 300–500 ng/μl) from different origins.

| Behaviours of the parasitoid *Cotesia flavipes* |  |
| --- | --- |
| Proteins tested | antennation + stinging attempt<br>(mean* ± SE, n=6) |
| A-amylase from *Aspergillus oryzae* | 43.3 ± 6.1bc |
| A-amylase from pig | 20.0 ± 7.3ab |
| A-amylase from *Drosophila melanogaster* | 70.0 ± 6.8d |
| A-amylase from *Chilo suppressalis* | 60.0 ± 7.3cd |
| Inactive α-amylase from *Drosophila melanogaster* | 53.3 ± 6.7cd |
| BSA | 0a |

\*Means with different letter are significant (q-value <0.05; pairwise Wilcoxon's rank sum test, q-value = FDR corrected p-value).

The α-amylases of both *D. melanogaster* and *C. suppressalis* induced action potentials from the gustatory neurons of sensilla chaetica located at the tip of antennae of *C. flavipes* females (Figs. 4 and 5); they were however weaker than those induced by the oral secretions of *C. partellus*. BSA induced action potentials equivalent to the control solution.

**Fig. 4.**
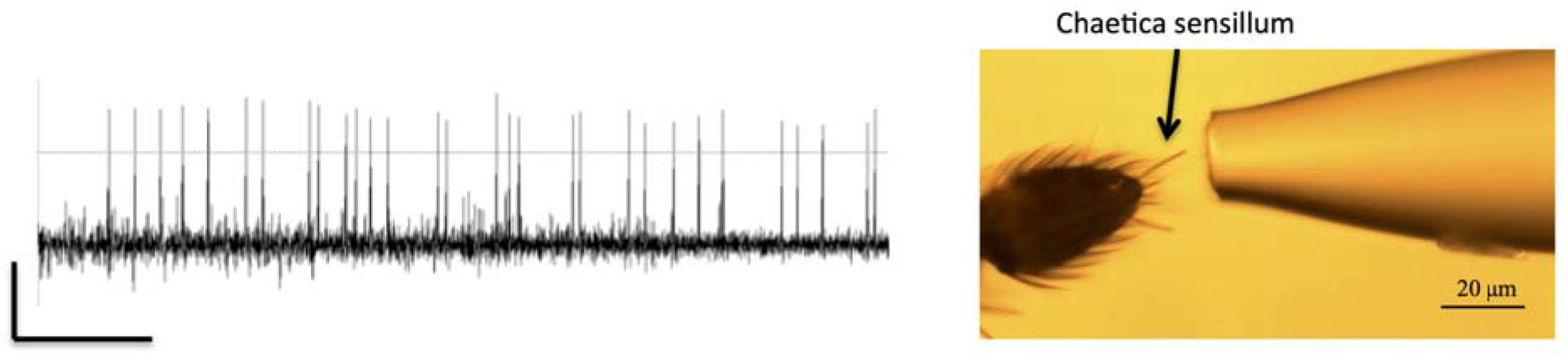
Left: 2 s chemosensory recording is displayed showing the response of a chaetica sensillum at the tip of *Cotesia flavipes* antennal female to α-amylase of *Drosophila melanogaster* (at 300 ng/μl). Vertical bar: 2 mV; horizontal bar: 200ms. Right: Photo of the tip of an antenna stimulated by a capillary electrode.

**Fig. 5.**
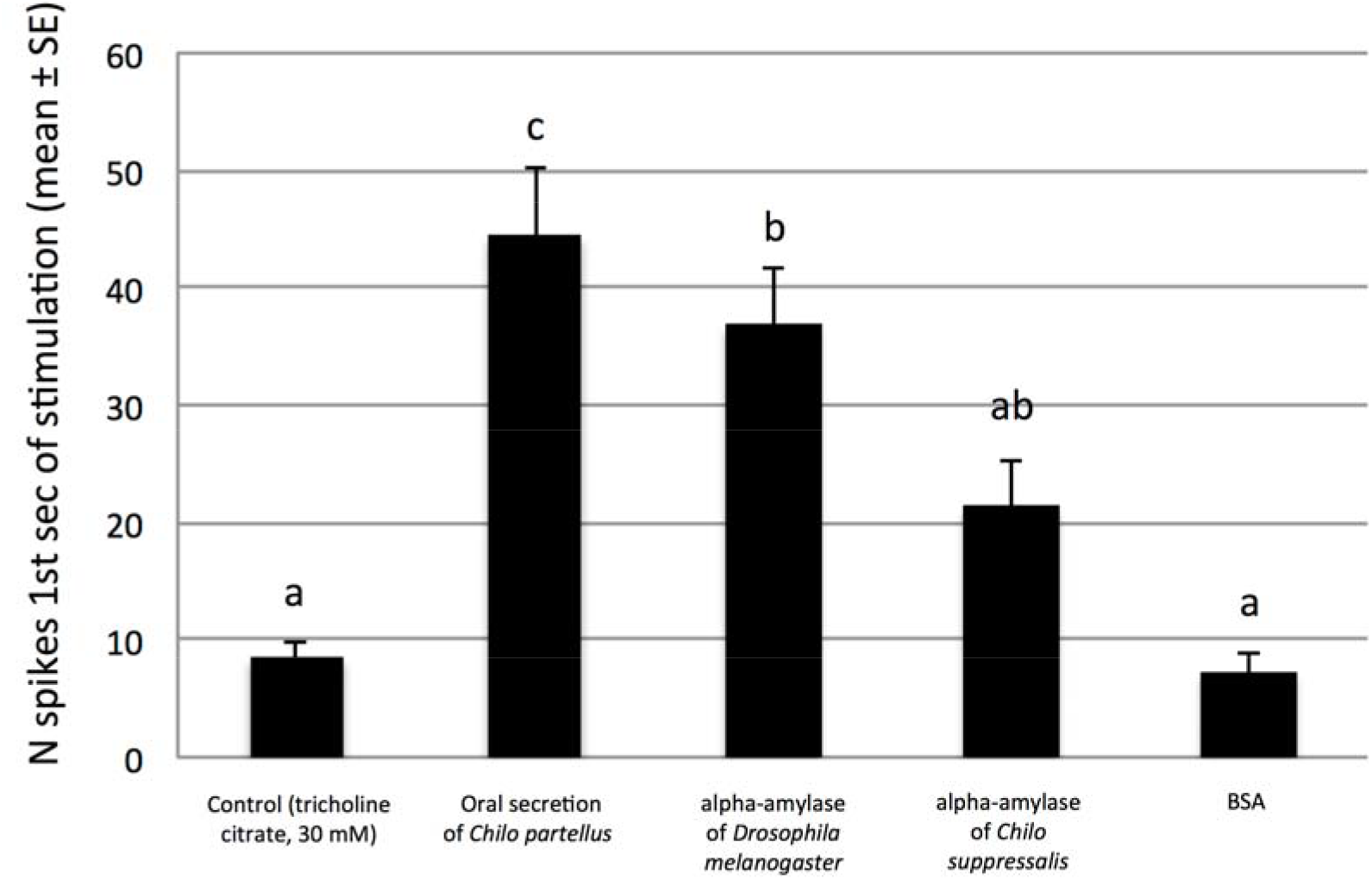
Electrophysiological responses of *Cotesia flavipes* females to oral secretion of *Chilo partellus* and to different proteins (at 300 ng/μl). The recordings were made on sensilla chaetica located at the apical antennal segments of *Cotesia flavipes* females. Each bar represents the mean (± SE, n=10) number of action potentials during the first second of stimulation. Different letters capping the bars indicate significant differences (P < 0.05) among mean responses elicited by the different stimuli (one-way ANOVA, Tukey’s contrasts test).

## DISCUSSION

In the current study, a compound involved in host acceptance for oviposition by the wasp *C. flavipes* isolated from the oral secretion of the larval host *C. partellus* was identified as an α-amylase. In *Pieris brassicae* (L) (Lepidoptera: Pieridae) larvae, the β-glucosidases of the oral secretion causes the release of VOCs from Brassicaceae plants that attract parasitoids (Mattiacci et al., 1995). Similarly, volicitin [N-(17-hydroxylinolenoyl)-L-glutamine] a compound present in the oral secretion of *Spodoptera* sp. (Lepidoptera: Noctuidae) induces the release of maize VOCs that attract parasitoids (Turlings et al., 1990; Alborn et al., 1997). In our study, although we have not tested yet if this enzyme induces the release of VOCs that can attract parasitoids, direct perception of the α-amylase upon contact elicits the antennation and stinging attempt behaviors of the parasitoid.

Although polypeptides and proteins have previously been reported as chemical signals in the host selection process by hymenopteran parasitoids (Weseloh, 1977; Bénédet et al., 1999; Gauthier et al., 2004), the definitive identification of such protein or polypeptide has never been achieved.

α-amylases are among the important classes of digestive enzymes used by the insects to hydrolyze starch to oligosaccharides in various plant tissues; thus they play a critical role in insect survival by providing energy (Franco et al., 2000). They have been found in several insect orders such as Orthoptera, Hemiptera, Heteroptera, Hymenoptera, Diptera, Lepidoptera and Coleoptera (Kaur et al., 2014). In Lepidoptera, α-amylases have variable molecular weights depending on the species (Sharifloo et al., 2016), which is unexpected since all known sequences of insect amylases predict roughly the same weight as those of *Drosophila melanogaster*, i.e. ≈ 50 kDa (Boer and Hickey, 1986; Titarenko and Chrispeels, 2000; Maczkowiak and Da Lage, 2006; Pytelkova et al., 2009; Bezerra et al., 2014; Channale et al., 2016).

The α-amylases tested in our study had a similar molecular weight as those of *D. melanogaster* (51 kDa) (C. *suppressalis:* ≈ 50 kDa, *A. oryzae*: 51 kDa and pig: 50 kDa). Interestingly, all these α-amylases induced behavioral responses of *C. flavipes*, suggesting that the size of the α-amylase is involved. However, a different protein such as BSA with a similar molecular weight was not inducing any behavioral response suggesting that the conformation of the protein rather than its weight is involved in host acceptance for oviposition behavior of the parasitoid. In fact, an inactive α-amylase of *D. melanogaster* (with a similar conformation of the active α-amylase) was still inducing behavioral responses of *C. flavipes*. This indicates that it is the conformation of the α-amylase rather than its catalytic site that induces this activity, and suggests that *C. flavipes* can perceive the α-amylase through its sensorial equipment.

Obonyo et al. (2010a) observed that female parasitoids (including *C. flavipes*) use the tip of their antennae to recognize and accept their host larvae for oviposition. They identified on the last antennal segment the presence of uniporous sensilla chaetica known to have gustatory functions in insects (Obonyo et al., 2011). Our study confirms that these sensilla chaetica are involved in the perception of non-volatile host cues as already shown by Iacovone et al. (2016) for the egg parasitoid, *Trissolcus brochymenae* Ashmead (Hymenoptera: Platygastridae). Gustation in insects is known to be influenced by small compounds such as sugars, free amino acids, water-soluble alkaloids (see Thiéry et al. [2013] for review), but the present findings demonstrate that it can also be elicited by larger molecular weight compounds such as proteins. In addition, as no action potential was generated by α-amylase from gustatory neurons of antennal sensilla chaetica of *C. flavipes* males (data not shown), such gustatory perception of α-amylase is most likely linked to host acceptance for oviposition behavior in *C. flavipes* females.

The implication of α-amylase in host recognition and thus selection for oviposition by the parasitoid implies a stable relationship between α-amylase variability among host larvae species and host specificity. In the last decade, it was observed that the diversity of Lepidoptera stemborers in Africa is considerably higher than described earlier (Le Ru et al., 2006a; 2006b) and that most of these stemborers are specialists (monophagous, oligophagous), exhibiting a strong host plant conservatism (Le Ru *et al*., 2006a; 2006b; Ong’amo *et al*., 2006a; 2006b; Otieno *et al*., 2006). In parallel, Mailafiya et al. (2009) found a higher diversity of the associated parasitoids than previously thought among *Busseola* spp. and *Chilo* spp. host genera, with an apparent strong host insect conservatism. The sequences of α-amylase gene (*Amy*) of a number of animals show a high level of protein variability (Da Lage et al., 2002). Therefore, the diversity of α-amylase proteins and of the corresponding *Amy* genes family may have adaptive or functional significance, for example, in the diversity of stem borers – parasitoids interactions. In fact, we observed a clearer and stronger behavioral response of *C. flavipes* with the oral secretion of *C. partellus* containing the genuine α-amylase than with all the other tested amylases.

A question therefore arises on how the parasitoids access α-amylase in nature? In fact, Lepidoptera stemborers larvae spend their life and feed inside plant stems. Before it enters into feeding tunnel of the host larvae, the wasp makes first contact with the fecal pellets left by the larvae pushed outside of the stem. Although, these pellets do not induce oviposition, they act as a marker of the status of the larva inside the stem tunnel as being host or non-host (Obonyo et al., 2010b) and if the host is still actively feeding or not. However, only when the parasitoid is in contact with the host body, it is able to recognize and accept it for oviposition (Obonyo et al., 2010a; 2010b). It is during this final step that the parasitoid can access the stimulatory compounds present on the body of the larvae deposited by their feeding activity. These stimulatory compounds need to give quick and appropriate information to the parasitoid on the suitability of the larva (both host and health status) because host larvae often bite the attacking wasps, causing a 50% mortality risk (Takasu and Overholt, 1997). The high selection pressure due to the high mortality at oviposition should favor wasps that are able to recognize their hosts with minimal risk of injury (Ward, 1992). Among the stimulatory compounds, this study shows α-amylases as good candidates for an evolutionary solution to host selection in parasitoids, opening new routes of investigation in hosts-parasitoids interactions.

## Acknowledgements

The authors whish to thank the University of Nairobi and *icipe* capacity building for hosting the PhD student. Thanks are given to Julius Obonyo and Peter Malusi for rearing the host insects and the parasitoids; and to Peter Ahuya and Kevin Sambai for their technical assistance in collecting the oral secretions and some behavioural assays, all from *icipe*. Thanks are also given to Fritz Schulthess for his review of the manuscript.

## Competing interests

The authors declare no competing or financial interests.

## Author contributions

P.-A.C. designed and supervised the study. G.B. carried out all the behavioral experiments, the electrophoresis and the isolation of proteins for identification. J.-L.D.L. synthesized the α-amylases from *Drosophila melanogaster* and *Chilo suppressalis* as well as the inactive α-amylase from *D. melanogaster*. J.-L.D.L. and C.C.-D. realized the western blot analysis. C.-M. M. performed the electrophysiological experiments. F.M.-P. supervised the electrophysiological experiments. M.Z. and T.B. realized the protein identification by LC-MS/MS. G.J., E.M. participated to the supervision of the study. G.B., B.L.R., L.K.-A., G.J., E.M. and P.-A.C. prepared the manuscript. L.K.-A. coordinated the research program hosting this work.

## Funding

This work was supported by IRD and ANR ABC PaPoGen for funding the research as well as DAAD for funding the PhD fellowship under the grant number 91560009.

## Supplementary information

**Table S1.** Results of proteins and peptides obtained by X!Tandem as well as proteins and peptides obtained by *de novo* (see attached excel table).

**Figure S1.**
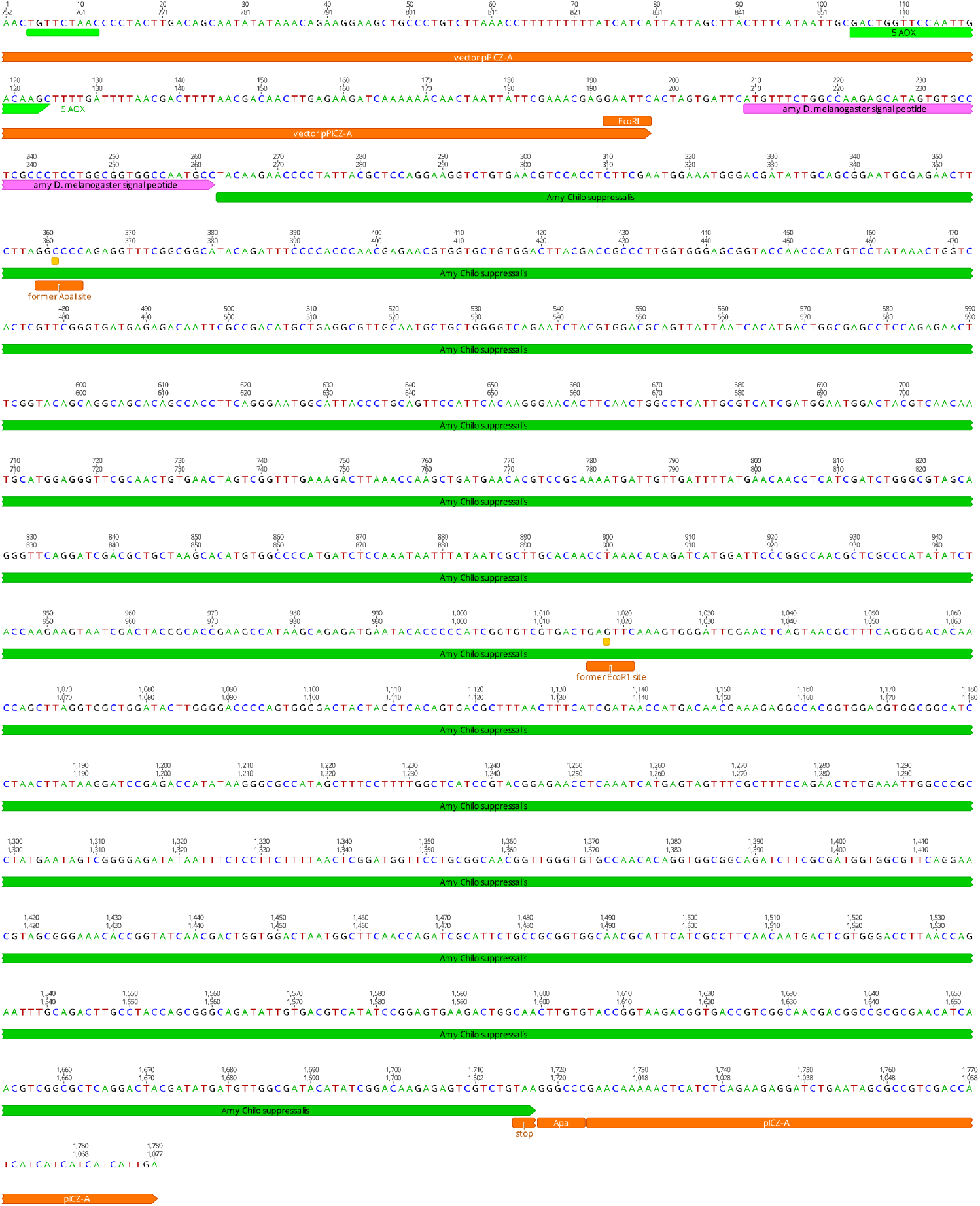
Map and sequence of the *Chilo suppressalis* 108827 amylase gene construct in the pPICZ-A expression vector (Invitrogen). The original signal peptide was replaced by the one of *Drosophila melanogaster* amylase. Two restriction sites were destroyed in the sequence to allow the use of those sites as cloning sites.

